# Eukaryotic stress induced mutagenesis is limited by a local control of Translesion Synthesis

**DOI:** 10.1101/2021.07.02.450853

**Authors:** Katarzyna H. Masłowska, Florencia Villafañez, Luisa Laureti, Shigenori Iwai, Vincent Pagès

## Abstract

The DNA Damage Response (DDR) preserves the genetic integrity of the cell by sensing and repairing damages after a genotoxic stress. Translesion Synthesis (TLS), an error-prone DNA damage tolerance pathway, is controlled by PCNA ubiquitination. In this report, we raise the question whether TLS is controlled locally, or globally. Using a recently developed method that allows to follow the bypass of a single lesion inserted into the yeast genome, we show that: i) TLS is controlled locally at each individual lesion by PCNA ubiquitination, ii) a single lesion is enough to induce PCNA ubiquitination, and iii) PCNA ubiquitination is an imperative requirement for TLS to occur. More importantly, we show that global PCNA ubiquitination that follows a genotoxic stress does not increase TLS at individual lesions. We conclude that unlike the SOS response in bacteria, the eukaryotic DDR does not promote TLS and mutagenesis.

## INTRODUCTION

Accurate DNA replication is essential for genome stability. Since DNA is constantly insulted by endogenous and exogenous DNA-damaging agents, organisms have evolved several mechanisms to deal with DNA damage. The DNA damage response (DDR) includes cell cycle arrest, lesion repair, and lesion tolerance (1). Numerous repair systems remove various modifications from DNA in an error-free manner (2). However, despite their efficient action, it is inevitable that some lesions might be present during replication. Most DNA damage impedes DNA synthesis by high-fidelity replicative DNA polymerases. Therefore, to complete replication and maintain cell survival in the presence of residual DNA damage, cells have evolved two lesion tolerance mechanisms: Damage Avoidance (DA) and Translesion Synthesis (TLS). Damage avoidance (also named strand switch, copy choice or homology directed gap repair) is a pathway relying on the information of the newly replicated sister chromatid to circumvent the lesion in an error-free manner ((3) (4), reviewed in (5)). Translesion synthesis is a potentially mutagenic pathway that employs specialized DNA polymerases able to insert a nucleotide directly opposite the lesion (reviewed in (6) and (7)).

How the DDR controls mutagenesis has been widely studied in prokaryotic cells: the SOS response, by increasing the expression level of TLS polymerases in response to a genotoxic stress, greatly contributes to mutagenesis and therefore to the adaptive response to environmental stress (8). A good example of this phenomenon is the importance of the SOS response in resistance to antibiotics (9). Moreover, experiments involving the study of a single lesion have shown that the pre-induction of the SOS system by a genotoxic stress greatly increases the level of TLS and mutagenesis at the studied lesion (10) (11). Hence, in bacteria, two factors contribute to the increase in mutagenesis in response to a genotoxic stress: i) the number of lesions (the higher the number of lesions, the higher the probability to generate a mutation); ii) the increased level of TLS polymerases in response to the induction of SOS (the more TLS polymerases, the higher probability to bypass the lesion by TLS). Thus, the SOS response is a global response that favors TLS and mutagenesis.

The eukaryotic DDR includes mostly posttranslational modifications such as phosphorylation and ubiquitination (12). Damage-induced transcriptional regulation is less common but has also been reported (13). PCNA ubiquitination regulates lesion tolerance in response to DNA damage. After the replication fork encounters a DNA lesion, PCNA stalls at the lesion site, and single-stranded DNA (ssDNA) is generated downstream of the lesion. The formation of RPA protein-coated ssDNA leads to the recruitment of the Rad6-Rad18 complex and the subsequent mono-ubiquitination of PCNA at lysine K164 (14) (15) (16). This mono-ubiquitination can be further extended by Rad5 and Ubc13-Mms2, through the formation of K63-linked ubiquitin chains (17) (18). It is well established that in eukaryotic cells, PCNA mono-ubiquitination stimulates TLS, while PCNA polyubiquitination triggers DA (reviewed in (19)).

Forward mutagenesis assays have shown that the mutation frequency rapidly increases with the amount of genotoxic stress inflicted to the cell. While it is expected that the level of mutagenesis increases with the number of lesions (more lesions lead to more mutations), it is not known if an additional regulatory mechanism also contributes to the increase in mutagenesis in eukaryotic cells. Traditional bulk approaches do not allow to determine if the mutagenesis level is solely correlated to the number of lesions, or if a more global DNA damage response also favors mutagenesis.

As PCNA ubiquitination controls TLS, and PCNA ubiquitination increases in response to genotoxic stress, it appears intuitive that such global response exists: the more PCNA is mono-ubiquitinated, the more TLS will be used by the cell. However, the existence of such global response has never been demonstrated.

In this study, we set out to determine whether the level of TLS is regulated solely at the local level, or if the amount of damage present in the cell could favor TLS in a more global manner. We have recently devised a method to introduce a single lesion in the genome of a yeast cell (Figure 1A) (20). Such approach allows to dissect the regulation of the tolerance mechanisms in different genetic backgrounds as well as in different conditions of external stress for the cell. We have used our assay to determine whether the ratio between TLS and DA at the level of a single lesion is modulated by increasing level of genotoxic stress resulting in global DNA damage response and abundant PCNA ubiquitination.

**Figure 1:**
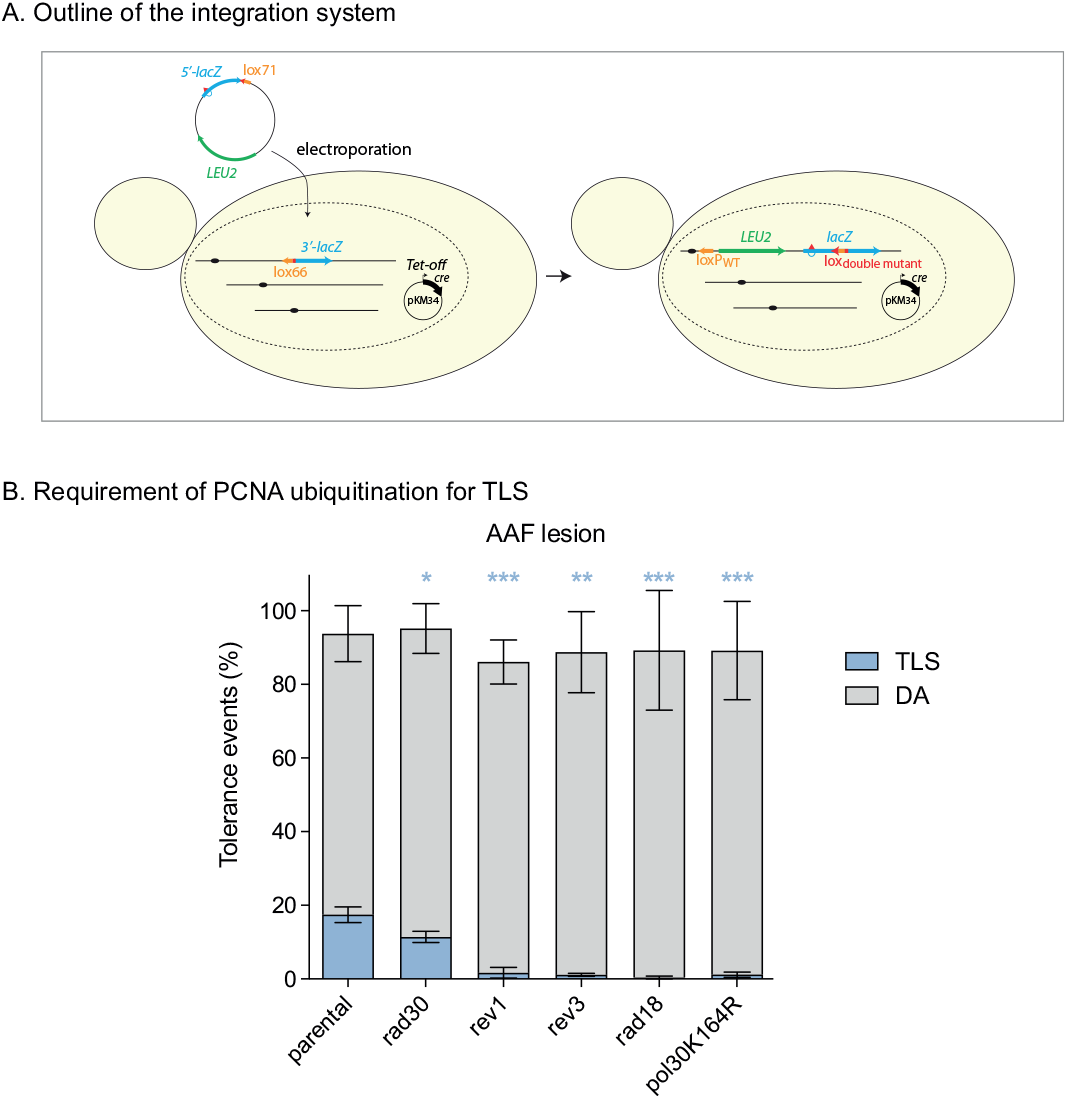
Requirement of PCNA ubiquitination for TLS. (A) Outline of the integration system: A non-replicative plasmid containing a single lesion is integrated into a yeast chromosome using a Cre/lox site-specific recombination. The integrative vector carrying a selection marker (LEU2) and the 5′-end of the lacZ reporter gene containing a single lesion is introduced into a specific locus of the chromosome with the 3′-end of lacZ. The precise integration of the plasmid DNA into the chromosome restores a functional lacZ gene, enabling the phenotypical detection of TLS and DA events (as blue and white colonies on X-gal indicator media). (B) Bypass of the N2dG-AAF lesion in strains deficient in TLS polymerases (Rev1, Pol η/ rad30, Pol ζ/rev3) and strains deficient in PCNA ubiquitination (rad18 and pol30K164R mutant).

## RESULTS

### PCNA ubiquitination occurs locally at a single lesion

The assay we have previously described (20) is based on a non-replicative plasmid containing a single lesion which is stably integrated into a specific site of the yeast genome. The precise integration of the plasmid DNA into the chromosome restores a functional lacZ gene, enabling the phenotypical detection of TLS and DA events (as blue and white colonies on X-gal indicator media). In order to study tolerance events, we inactivated the following repair mechanisms: nucleotide excision repair (rad14), photolyase (phr1), and mismatch repair system (msh2). Tolerance events are calculated as the ratio of colonies resulting from the integration of the damaged vector versus the non-damaged vector.

Using this method, we have introduced a N2dG-AAF (N2-dG-Acetylaminofluorene) adduct in the genome of cells deficient for repair mechanisms, but proficient for lesion tolerance. 2-Acetylaminofluorene is a potent carcinogen known to produce liver tumors in rats (26). We observe for this lesion a TLS level of 18% that relies almost exclusively on the TLS polymerases Rev1 and Pol ζ (Figure 1B). We also observed a reduction of TLS in the rad30 mutant showing that Pol η is also involved in the bypass of this lesion as previously shown (27). Indeed, it has been previously suggested that Pol η can participate to the insertion step at the N2dG-AAF lesion, while Pol ζ is required for the extension step (27).

We can note that the level of TLS is highly dependent on the type of lesion as we previously measured 55% of TLS for the cis-syn TT dimer lesion (cyclobutane pyrimidine dimer) and 5% of TLS for the (6-4)TT photoproduct lesion (thymine-thymine pyrimidine(6-4)pyrimidone photoproduct) (20). Replication through all of these lesions relies exclusively on TLS polymerases as the inactivation of rad30, rev3 and/or rev1 abolishes TLS (20).

We then introduced the N2dG-AAF lesion in cells that were unable to ubiquitinylate PCNA, either in rad18 strains, or strains where the lysine 164 of PCNA was mutated into an arginine (pol30K164R) (Figure 1B). In both conditions, TLS was almost completely abolished which is consistent with the current view of PCNA ubiquitination promoting TLS. The same results were previously observed for the cis-syn TT dimer and (6-4)TT photoproduct lesions (20). Therefore, PCNA ubiquitination appears as an imperative requirement for TLS to occur since almost no TLS can be achieved in its absence. Given that the lesions were introduced in the cells in the absence of any other DNA damaging treatment, the presence of the single lesion is enough to generate the signal required to trigger Rad6-Rad18-mediated PCNA ubiquitination. Hence, these data indicate that PCNA ubiquitination occurs locally, and that the single patch of ssDNA generated at one lesion is enough to recruit the Rad6-Rad18 complex that will ubiquitinate PCNA allowing TLS to occur at the lesion. We can note here that the DA level remains high despite the absence of PCNA ubiquitination. These events can be attributed to the “salvage recombination pathway” that has previously been described (5). This pathway allows to bypass the lesion using homologous recombination, but occurs independently of Rad6-Rad18 (28).

We then wondered if in addition to this local regulation, a more global control over PCNA ubiquitination and TLS exists. In other words, can the substantial level of PCNA ubiquitination generated by a genotoxic stress increase the use of TLS at each lesion?

### Is TLS modulated by a global stress response?

To answer this question, we treated the cells with DNA damaging agents (either 4-NQO: 4-Nitroquinoline-1-oxide (29), or UV irradiation) prior to the integration of three types of lesions (cis-syn TT dimer, (6-4)TT photoproduct or N2dG-AAF). We used a dose that causes about 80% lethality. Such treatment is known to induce a strong DDR, which includes PCNA ubiquitination and Rad53 phosphorylation. Indeed, we showed by western blot that 4-NQO treatment leads to significant PCNA ubiquitination (Figure 2B) and Rad53 phosphorylation (Figure 2C). Similarly, we showed that UV-irradiation leads to PCNA ubiquitination (supplementary Figure 1). We also verified that the treatment to make the cells competent for electroporation and the electroporation itself were not inducing stress triggering PCNA ubiquitination. This confirms that the TLS data obtained previously (Figure 1) is independent of any genotoxic stress and only related to the single inserted lesion.

**Figure 2:**
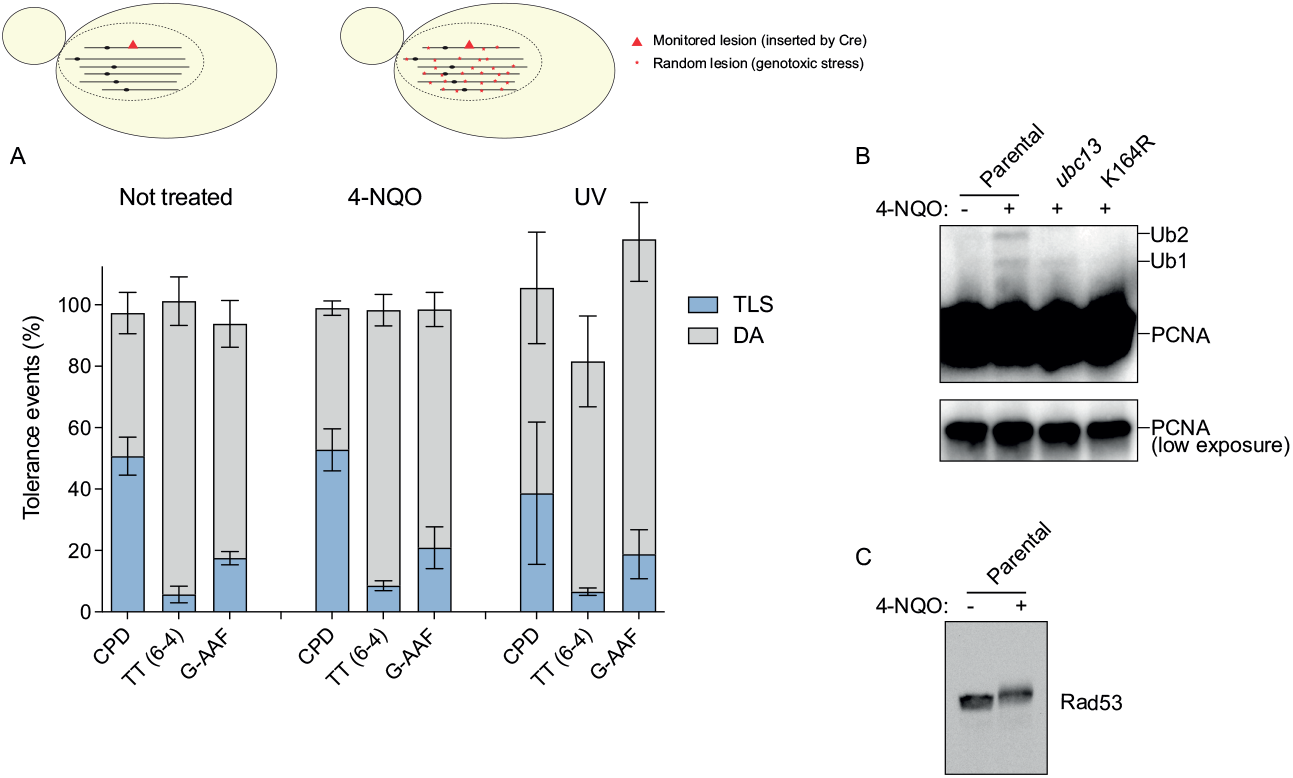
TLS is not modulated by a global stress response. (**A**) Partition of DNA Damage Tolerance events for different lesions (CPD, TT (6-4) and G-AAF) non-treated (left) and treated with 4-NQO or UV irradiation (right) prior to the integration. No significant difference was observed between the non-treated and treated conditions. (**B**) Western blot analysis revealing flag-tagged PCNA shows that the electroporation conditions do not induce significant level of PCNA ubiquitination. Upon treatment, two bands appear that correspond to mono- and bi-ubiquitination of PCNA. In the *pol30*-K164R mutant, these two bands are no longer present since the site of ubiquitination of PCNA (Lysine 164) is absent. In the *ubc13* mutant, the polyubiquitination band disappears, leaving only the monoubiquitinated version of PCNA. (C) Western blot analysis revealing Rad53 phosphorylation in response to 4-NQO treatment.

Having confirmed that PCNA is ubiquitinated in response to 4-NQO or UV treatment, we introduced a single lesion cis-syn TT dimer, (6-4)TT photoproduct or N2dG-AAF) after treating the cells with 4-NQO for 30 minutes, or after UV-irradiation. No increase in TLS was observed in cells pre-treated by 4-NQO as compared to the nontreated cells. The same results were obtained with UV-irradiated cells (Figure 2A).

It appears therefore that the substantial level of PCNA ubiquitination resulting from genotoxic stress does not modulate the bypass of individual lesions. This suggests that a preexisting pool of ubiquitinated PCNA does not affect the bypass of the newly appeared lesion. Similarly, the activation of the DDR (monitored by Rad53 phosphorylation) does not modify the level of TLS at the integrated lesion. This implies that TLS is controlled at a local level, rather than by a global DDR.

### Global vs. local response to DNA damage

To confirm this hypothesis, and to avoid any bias that could be introduced by our integration system, we compared UV-induced mutagenesis in bacterial and yeast strains (Figure 3). In short, cells were UV-irradiated at different UV doses and mutants were counted as Rifampicin resistant colonies for E. coli (30), or as Canavanine resistant colonies for S. cerevisiae (31).

**Figure 3:**
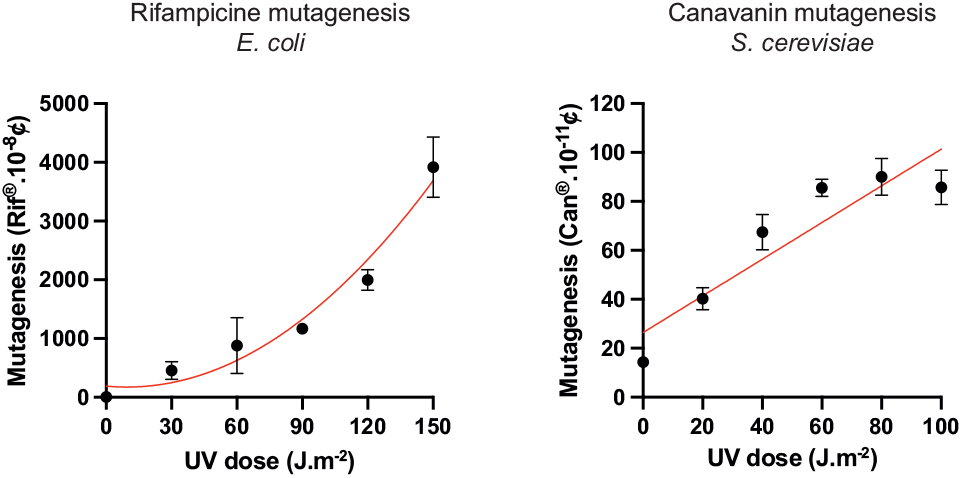
UV-induced mutagenesis in *E. coli* and *S. cerevisiae. E. coli* mutants were scored as rifampicin resistant colonies, *S. cerevisiae* as canavanine resistant colonies. Each point represents the average and standard deviation of at least three independent experiments. The curve in red represents the fit following a quadratic equation for the Rif mutagenesis (R^2^ =0.93) or a linear fit for the Can mutagenesis (R^2^ =0.82).

In E. coli, we observe a quadratic correlation between the UV dose and mutagenesis. Indeed, the number of mutations increases with both the number of lesions, and with the probability of a lesion to turn into a mutation (that also increases with the amount of lesions that induce the SOS response). Consequently, the number of mutations increases with the square of the number of lesions (quadratic correlation). This reflects the global effect of the SOS response.

On the other hand, in yeast we observe a linear correlation that even tends to reach a plateau for high doses. This shows that the genotoxic stress does not favor TLS, and mutagenesis only increases with the number of lesions. This reflects the absence of global regulation of TLS in this organism.

## DISCUSSION

Our data show that a single lesion is enough to induce PCNA ubiquitination locally, and that PCNA ubiquitination is an imperative requirement for TLS to occur. More importantly, we show that a genotoxic stress leading to a significant level of PCNA ubiquitination and to the activation of the DNA damage response (Rad53 phosphorylation) has no effect on the level of TLS at individual lesions. We conclude from this data that the eukaryotic DNA damage response does not favor the increase of TLS and mutagenesis. This observation was confirmed by UV induced mutagenesis experiments showing that the mutagenic response increases only linearly with the level of damage in yeast. This response is quite different from the one in E. coli where it has previously been shown that the level of TLS at individual lesion increases upon UV-irradiation (10), and where the mutagenic response shows a quadratic correlation with the UV dose.

The fact that the increased level of PCNA ubiquitination does not result in an increase of TLS or mutagenesis suggests that ubiquitinated PCNA is not recycled and does not stimulate TLS at other lesion sites. This implies that upon completion of a TLS patch, PCNA is either rapidly deubiquitinated, or that PCNA that is unloaded is deubiquitinated before it is recycled at another replication fork, so it will not favor TLS at another lesion site. This rapid deubiquitination could explain the low level of PCNA ubiquitination generally observed upon genotoxic stress (this study and (16)). Indeed, the level of Ub-PCNA increases when the deubiquitylases are inactivated (32).

This also implies that de novo ubiquitination occurs at each encounter with a lesion. This model is compatible with previous in vitro data that showed that PCNA remains at the lesion terminus where RPA coated ssDNA prevents its diffusion (33). In this model, a new PCNA is loaded downstream of the lesion allowing the replication to go on, while the PCNA that is maintained at the lesion site can be ubiquitinated and allows TLS to perform the gap filling reaction. Once the gap filling reaction is achieved, ubiquitinated PCNA is unloaded and will not contribute to TLS at another site.

Other events of the DDR response could affect the level of TLS besides PCNA ubiquitination. For instance, TLS polymerase Rev1 has been reported to be phosphorylated in response to a genotoxic stress (34), which was demonstrated to be important for DNA damage bypass (35). It appears from our data that such modifications are not playing a global role, but again have most likely a local effect at each lesion.

Taken together, our data show that the regulation of TLS occurs locally, and that in eukaryotic cells, there is no global response capable of increasing the level of TLS and mutagenesis in response to a genotoxic stress. The level of mutagenesis depends solely on the number of lesions present in the genome (following a linear correlation). Similar mutagenesis assays in human cells have shown the same linear response (36, 37). Unlike bacteria that show a quadratic increase in mutagenesis, allowing them to rapidly mutate and adapt to a stressful environment, eukaryotic cells seem to have evolved a more controlled mutagenic response to the environmental stress.

## MATERIAL AND METHODS

### Strains and media

All strains used in the present study are derivative of strain EMY74.7 (21) (MATa his3-Δ1 leu2-3,112 trp1-Δ ura3-Δ met25-Δ phr1-Δ rad14-Δ msh2Δ::hisG). Gene disruptions were achieved using PCR-mediated seamless gene deletion (22) or URAblaster (23) techniques. All strains used in the study are listed in Table 1.

**Table 1:** Strains used i, the study

| Strain | Relevant Genotype |
| --- | --- |
|  | (All yeast strains are: *MATa his3-Δ1 leu2-3,112 trp1-Δ ura3-Δ met25-Δ rad14-Δ phr1-Δ msh2Δ::hisG*) |
| EVP161 | *E. coli* MG1655 *phrB* ::kan |
| SC22 | *rad14 phr1* |
| SC53 | VI(167260-167265)::(*lox66-3'lacZ-MET25/lag*) |
| SC55 | VI(167260-167265)::(*lox66-3'lacZ-MET25/lead*) |
| SC82 | *rev1-Δ* VI(167260-167265)::(*lox66-3'lacZ-MET25/lag*) |
| SC83 | *rev1-Δ* VI(167260-167265)::(*lox66-3'lacZ-MET25/lead*) |
| SC86 | *rad30Δ::hisG* VI(167260-167265)::(*lox66-3'lacZ-MET25/lag*) |
| SC87 | *rad30Δ::hisG* VI(167260-167265)::(*lox66-3'lacZ-MET25/lead*) |
| SC181 | *rev3Δ::hisG* VI(167260-167265)::(*lox66-3'lacZ-MET25/lag*) |
| SC182 | *rev3Δ::hisG* VI(167260-167265)::(*lox66-3'lacZ-MET25/lead*) |
| SC203 | *rad18Δ::hisG* VI(167260-167265)::(*lox66-3'lacZ-MET25/lag*) |
| SC206 | *rad18Δ::hisG* III(75494-75499)::(*lox66-3'lacZ-MET25/lead*) |
| SC236 | *pol30-K164R* VI(167260-167265)::(*lox66-3'lacZ-MET25/lag*) |
| SC237 | *pol30-K164R* VI(167260-167265)::(*lox66-3'lacZ-MET25/lead*) |
| SC533 | *pol30::3FLAG-kan* VI (167260-167265)::(*lox66-3'lacZ-MET25/lag*) |
| SC534 | *pol30::3FLAG-kan* VI (167260-167265)::(*lox66-3'lacZ-MET25/lead*) |
| SC535 | *ubc13Δ* *Pol30::3FLAG-kan* VI (167260-167265)::(*lox66-3'lacZ-MET25/lag*) |
| SC537 | *pol30-K164R::3FLAG-kan* VI (167260-167265)::(*lox66-3'lacZ-MET25/lag*) |

### Integration system

Integration of plasmids carrying cis-syn TT dimer / 6-4 (TT) / N2dG-AAF lesions (or control plasmids without lesion) and result analysis was performed as previously described (20). In experiments where cells were treated with 4-NQO, the overnight culture was inoculated into 100 ml of yeast extract/peptone/dextrose medium (YPD) per integrated lesion to reach OD600 = 0.3, and incubated at 30°C with shaking until OD600= 0.8. After the addition of 150 ng/ml of 4-NQO, cultures were incubated for 30 more minutes. 4-NQO was inactivated by adding an equal volume of 10% sodium thiosulfate, and then cells were further washed and processed the same way as untreated cultures. For UV treatment, the overnight culture was inoculated into 50 ml of YPD per integrated lesion to reach OD600 = 0.3, and incubated at 30°C with shaking until OD600=1.6. Cells were then harvested, resuspended in twice the initial volume of water, and treated with UV (4J.m-2) in Petri dishes (15 cm Æ, 25 ml/dish). Cells were further washed and processed the same way as untreated cultures.

All experiments were performed in triplicate or more. Only the N2dG-AAF lesion in the UV-induced condition was done in duplicate. Graphs and statistical analysis were performed using GraphPad Prism applying unpaired t-test. Bars represent the mean value ± s.d.

### Preparation of an oligonucleotide containing the (6–4) photoproduct

A 13-mer oligonucleotide, d(GCAAGTTAACACG), purchased from Tsukuba Oligo Service (150 nmol) was dissolved in water (7.5 mL), and after a nitrogen purge for 5 min, the solution was heated at 75°C for 5 min and cooled in ice. This solution was poured into a petri dish with a diameter of 8.5 cm and irradiated for 20 min on a Spectrolinker XL-1500 UV Crosslinker (Spectronics Corp.) equipped with six 15 W germicidal lamps (254nm, 2.2 mW.cm-2). The reaction mixture was analyzed by reversed-phase HPLC using a Waters μBondasphere C18 5μm 300Å column (3.9’ 150 mm) at 50°C with an acetonitrile gradient of 6 to 10% during 20 min in 0.1 M triethylammonium acetate (pH 7.0). The elution was monitored by using a Waters 2998 photodiode array detector, and the product with absorption at 325 nm (which is characteristic of the (6-4) photoproduct) was isolated by repeating injection of 500 μL. After concentration of the combined eluates, impurities detected by the second HPLC analysis using a GL Sciences Inertsil ODS-3 5μm column (4.6’ 250 mm) with an acetonitrile gradient of 8 to 11% were removed. The final eluate (chromatogram shown in Supplementary Figure 2) was concentrated and co-evaporated with water. The yield determined from the absorbance at 260 nm was 4.5 nmol. This process is similar to the one previously described by LeClerc et al. (24).

### Immunoprecipitation and Western-Blot

Cells expressing FLAG-tagged PCNA were grown and processed as for the integration of the plasmid with a lesion. After electroporation cells were left to recover during 30 min, harvested, washed with buffer containing glycerol (1x PBS, 10% glycerol, and 1 mM EDTA), and frozen at −80°C. For protein extraction cells were resuspended in protein extraction buffer (1xPBS, 10% glycerol, 1mM EDTA, 1mM PMSF, 1x Roche cOmplete™Protease Inhibitor Cocktail), and lysed in a bead beater. Lysate was clarified by centrifugation, and total protein concentration was determined using Qubit Fluorometer according to manufacturer’s protocol (ThermoFisher). Crude protein extract was incubated with Anti-FLAG^®^ M2 Magnetic Beads (Merck) overnight, in an IP buffer (1xPBS, 5% glycerol, 0.5 mM EDTA, 1mM PMSF, 1x Roche cOmplete™Protease Inhibitor Cocktail, 2mM DTT). Proteins were resolved by SDS-PAGE (Bio-Rad 12% Mini-PROTEAN^®^ TGX™ Precast Protein Gels) and transferred onto PVDF membranes using Bio-Rad Trans-Blot Turbo Transfer System for Western blotting. Modified and non-modified PCNA was detected using ANTI-FLAG^®^ M2-Peroxidase (HRP) antibody (Merck).

For Rad53 detection, protein extracts for Western blot analysis were prepared by trichloroacetic acid (TCA) precipitation, as described previously (25). Proteins were resolved by SDS-PAGE (Bio-Rad 10% Mini-PROTEAN^®^ TGX™ Precast Protein Gels) and transferred onto PVDF membranes. Rad53, phosphorylated and not, was detected using anti-Rad53 polyclonal antibodies (Abcam ab104232).

### Rifampicin mutagenesis assay

E. coli strain EVP161 (MG1655 phrB::kan) was grown to exponential phase in LB media, centrifugated and resuspended in 10 mM MgSO4 before UV irradiation with different UV doses (0-150 J.m-2). An aliquot of cells was plated on LB to assess survival. Cells were then diluted 1/20 in LB and grown for 6h before plating on LB and LB + Rif 100μg/ ml. Colonies were counted after 24h on LB and 48h on LB+Rif.

### Canavanine mutagenesis assay

Mutagenesis experiments were performed as described in Unk and Daraba (2014). Briefly, yeast cultures were grown to saturation in YPD, harvested, resuspended in water (108 cells/ml), and sonicated to separate clumps of cells. Serial dilutions were plated on SD-agar plates containing canavanine for mutagenesis, and on SC-agar plates for survival. Plates were left to absorb the moisture, and irradiated with different doses of UV (0-100 J.m-2). Colonies were counted after 4 days of incubation.

## FUNDING

Fondation pour la Recherche Médicale: Equipe FRM - EQU201903007797

Fondation de France

