## Supplementary information for "Eukaryotic stress induced mutagenesis is limited by a local control of Translesion Synthesis"

___

Katarzyna H. Masłowska[1], Florencia Villafañez[1], Luisa Laureti[1], Shigenori Iwai[2], Vincent Pagès[1*]

[1]Cancer Research Center of Marseille: Team DNA Damage and Genome Instability | CNRS, Aix Marseille Univ, INSERM, Institut Paoli-Calmettes, Marseille, France
[2]Graduate School of Engineering Science, Osaka University, Osaka, Japan
[*]

___

**Supplementary data**

**Supplementary Figure 1:**

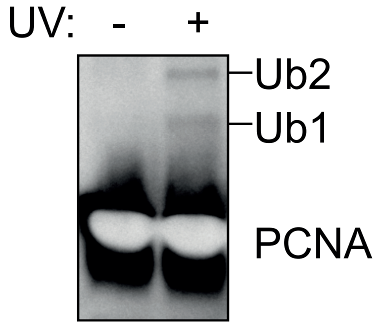


**Supplementary Figure 1:** Western-blot analysis for the UV irradiation condition revealing a significant increase in PCNA ubiquitination in the treated condition.

**Supplementary Figure 2:**

### HPLC control of the (6-4)TT photoproduct oligonucleotide

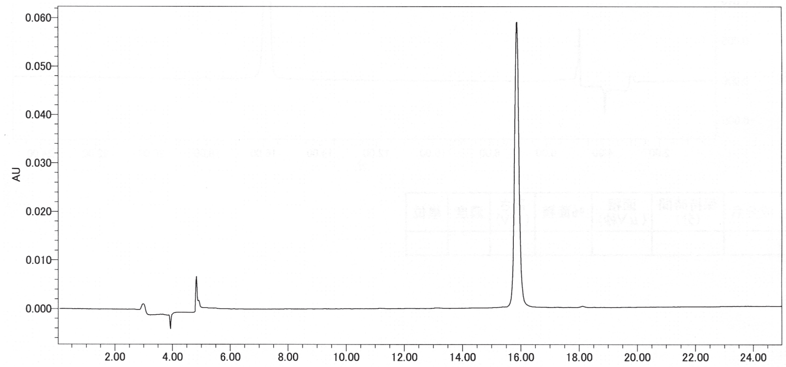


**Supplementary Figure 2:** Chromatogram showing the purity of the (6-4) TT photoproduct oligo following 2 chromatography columns (as describe in the method section).
